# An *Enterobacteriaceae* bloom in aging animals is restrained by the gut microbiome

**DOI:** 10.1101/2023.06.13.544815

**Authors:** Rebecca Choi, Rahul Bodkhe, Barbara Pees, Dan Kim, Maureen Berg, David Monnin, Juhyun Cho, Vivek Narayan, Ethan Deller, Michael Shapira

## Abstract

The gut microbiome plays important roles in host function and health. Core microbiomes have been described for different species, and imbalances in their composition, known as dysbiosis, are associated with pathology. Changes in the gut microbiome and dysbiosis are common in aging, possibly due to multi-tissue deterioration, which includes metabolic shifts, dysregulated immunity, and disrupted epithelial barriers. However, the characteristics of these changes, as reported in different studies, are varied and sometimes conflicting. Using clonal populations of *C. elegans* to highlight trends shared among individuals, and employing NextGen sequencing, CFU counts and fluorescent imaging to characterize age-dependent changes in worms raised in different microbial environments, we identified an *Enterobacteriaceae* bloom as a common denominator in aging animals. Experiments using *Enterobacter hormachei*, a representative commensal, suggested that the *Enterobacteriaceae* bloom was facilitated by a decline in Sma/BMP immune signaling in aging animals and demonstrated its detrimental potential for increasing susceptibility to infection. However, such detrimental effects were context-dependent, mitigated by competition with commensal communities, highlighting the latter as determinants of healthy versus unhealthy aging, depending on their ability to restrain opportunistic pathobionts.

## Introduction

Aging is a process of multi-tissue deterioration, including muscular atrophy, neurodegeneration, epithelial barrier disruption, immune dysregulation, and metabolic remodeling. Vulnerabilities and pathologies associated with this deterioration directly impact lifespan. In the case of the intestine, age-dependent impairments (immune, barrier, metabolic) further converge to alter the niche that is home to a complex community of microbes, the gut microbiome. However, how such changes affect the gut microbiome is not well understood.

The gut microbiome is increasingly appreciated for its contributions to host functions ^1–4^ and imbalances in its composition, or dysbiosis, are associated with pathology, in some cases (e.g. obesity) in a causative role ^5,6^. Age-dependent dysbiosis was described in flies, mice and humans and was suggested to negatively impact both barrier functions and immune fitness ^7–9^. Studies in human populations have shown that microbiomes of healthy octogenarians differed from those of unhealthy individuals of similar age ^10^. Other studies characterized the trajectory of microbiome changes through aging all the way to semi-supercentenarian (105-110 year old) ^11^, offering further insights into the relationship between age-dependent changes in microbiome composition and host health. Importantly, transplanting microbiomes from young mice to old reduced markers of aging and ameliorated health ^12,13^, and a similar transfer in killifish increased lifespan ^14^, demonstrating a causal role for age-dependent changes in microbiome composition in host aging. However, for the most part, such studies could not identify common trends in age-dependent changes that may offer points for intervention to ameliorate pathologies. Perhaps the sole exception is the identification of increased abundance of *Proteobacteria* (now re-named *Pseudomonadota*) in aging animals. Bacteria of this phylum are minor constituents of the vertebrate gut microbiome in young individuals, but increase in abundance during aging ^15–17^. *Pseudomonadota* comprise a larger part of the gut microbiome in invertebrates, and as seen in fruit flies further increase during aging ^7^. Whether this bloom is a universal signature of the aging gut microbiome, and what its significance may be, is yet unknown.

The nematode *Caenorhabditis elegans*, a useful model for aging research, is now gaining momentum as a model for microbiome research, offering the advantage of working with synchronized, initially germ-free, clonal populations, overcoming limitations of inter-individual variation common to vertebrate models to better discern shared patterns in microbiome composition ^18^. As a bacterivore, *C. elegans* ingests bacteria from its environment. While some bacteria are digested as food, others persist, giving rise to a characteristic gut microbiome that is surprisingly diverse, distinct from microbial communities in its respective environments, and similar in worms isolated from different geographical locations ^19–21^. As in other organisms, gut commensals were shown to provide diverse benefits to their host, including faster development and resistance to pathogens ^20–24^. The power of *C. elegans* as a genetic model further enabled identification of some of the genes, many of which are immune regulators ^25–27^, that help control commensals and their function and shape microbiome composition.

Here, we used *C. elegans* to characterize changes in gut microbiome composition during aging, identifying a bloom in bacteria of the *Enterobacteriaceae* family that was associated with an age-dependent decline in immune DBL-1/BMP signaling. The *Enterobacteriaceae* bloom was found to have the potential to be detrimental by increasing vulnerability to infection. However, competing commensals, or a diverse microbiome, were able to mitigate these detrimental effects. The results presented highlight an *Enterobacteriaceae* bloom as a hallmark of normal aging and suggest that the outcomes of this bloom are context-dependent, determined by the ability of the rest of the gut microbiome to restrain it, distinguishing between healthy and non-healthy aging.

## Results

### *C. elegans* aging involves an expansion in bacteria of the *Enterobacteriaceae* family

Worms continuously developing and aging in natural-like microcosm environments and analyzed by 16S next generation sequencing (NGS) showed gut microbiomes that were distinct from their environment, as previously described ^19^. The composition of these gut microbiomes changed as worms aged, but independently of bacterial environmental availability, which remained relatively constant during the experiment. (Fig. 1B, grazed soil). Most prominently, we observed an expansion in gut *Enterobacteriaceae*, particularly in post-reproductive worms (post-gravid, day five of adulthood). This increase could be due to ecological succession, shaped by interbacterial interactions. Alternatively, it could be determined by age-dependent changes in the gut niche. To distinguish between the two possibilities, we carried out microcosm experiments where worms of advancing age were exposed to the complex microcosm community for a fixed amount of time (Fig. 1A, C). While the initial soil microbiome in this experiment showed relatively higher microbial diversity compared to the experiment described in Fig. 1B, the *Enterobacteriaceae* expansion re-emerged, suggesting that this expansion was associated with age-dependent changes in the intestinal niche, rather than with the time of exposure, and highlighting this bloom as a potential hallmark of worm aging (Fig. 1C).

**Figure 1.**
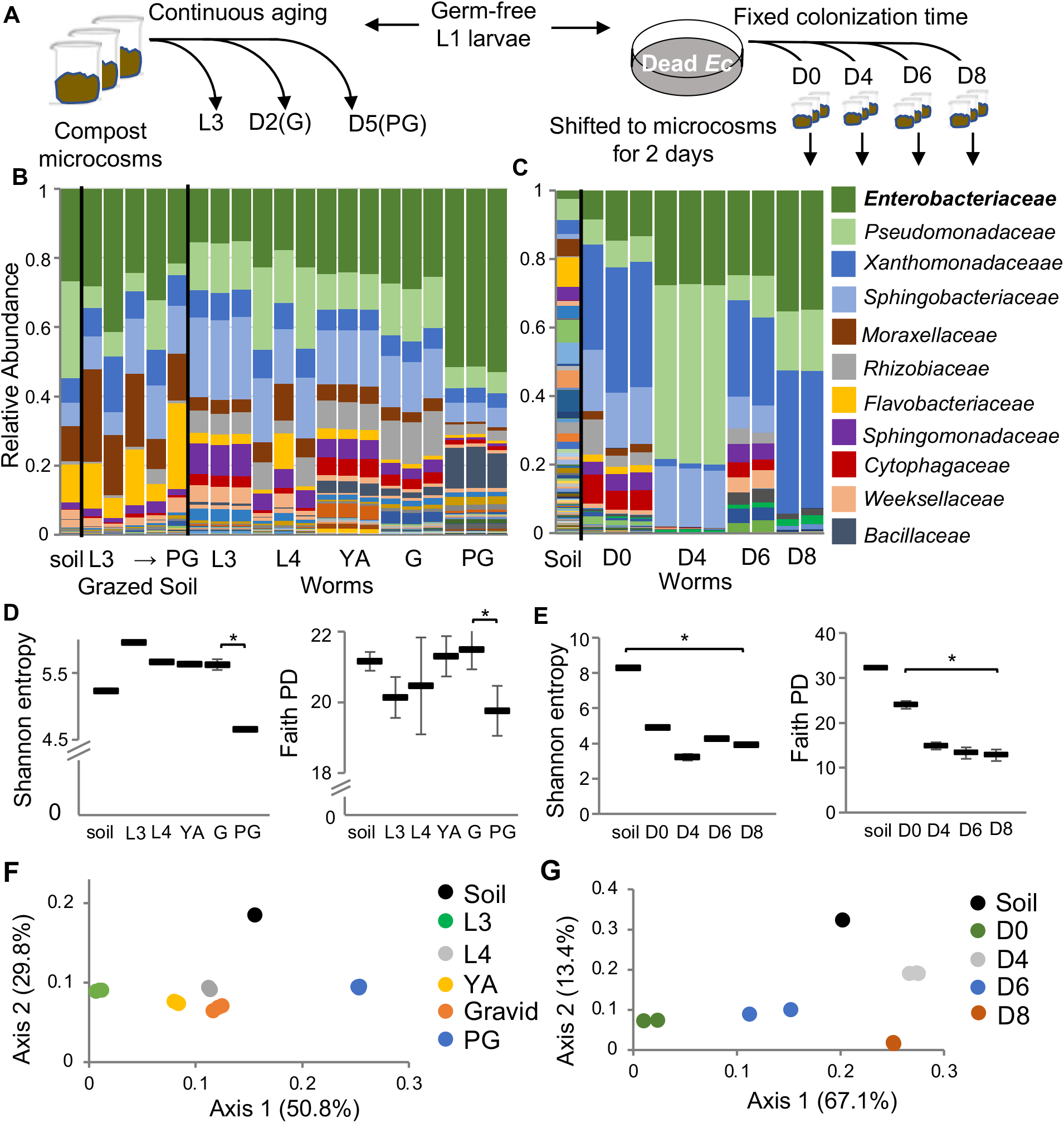
A gut *Enterobacteriaceae* bloom in worms aging in natural-like environments. **A**. Two sampling schemes for worm microbiome analysis during late development and early aging. **B**,**C**. Microbiome composition in worms raised continuously in microcosm environments (B) or in worms of advancing ages, shifted for two days to microcosm environments (C). L3, L4, larval stages; G, gravids (D2, second day of adulthood); PG, post-gravid (D5); D0 = early gravids. Bars represent microbiomes in microcosm environments or in the gut of worms raised in these microcosms (each bar represents a population of 100 worms). Taxa are shown at family level resolution. **D**,**E**. Alpha diversity represented by Shannon and Faith phylogeny indices; *, p<0.05, t-test. **F**,**G**. Principal Coordinate Analysis based on weighted UNIFRAC distances between microbiomes.

Bacterial diversity (alpha diversity) within the gut microbiome demonstrated an overall trend of decline during aging, which was more pronounced in the fixed-colonization-time aging experiment (Fig. 1E), but also seen in post-gravid worms in the continuous aging experiment (Fig. 1D). Declines were observed in worm microbiome diversity both with respect to species richness and evenness, represented by the Shannon Index, as well as with respect to phylogenetic diversity, represented by the Faith Index (Fig. 1D, E). Additionally, worm gut microbiomes of different ages differed from one another. Principal coordinate analysis (PCoA) based on weighted UNIFRAC distances showed that in both experiments worms of a specific age harbored similar gut microbiomes, which were distinct from worm microbiomes in other ages (Fig. 1F, G). This is in agreement with previous studies of the gut microbiome in aging mice ^28^. Together, these results support a role for the age-modified intestinal niche in driving age-dependent changes in microbiome composition, including a prominent expansion of *Enterobacteriaceae* as well as a general decline in bacterial diversity.

### An *Enterobacteriaceae* expansion is a recurring theme in aging starting from different initial conditions

Microcosm experiments provide natural-like microbial diversity, and coupled with NGS, offer a view free of culturing biases. However, to investigate the *Enterobacteriaceae* bloom in greater detail we turned to defined communities of characterized gut commensals. Two communities were used, each representing a slightly different paradigm: the publicly available CeMBio community, which includes two *Enterobacteriaceae* strains, *Enterobacter hormachei* strain (CEent1) and *Lelliottia amnigena* (JUB66), but is dominated by *Stenotrophomonas* and *Ochrobactrum*, ^29^ and SC20, which has a higher proportion of *Enterobacteriaceae* (eight out of 20 species), including CEent1 (Table 1). We used a CEent1-dsRed derivative included in both communities to enable imaging of gut colonization. In both, colonization with CEent1-dsRed increased with age, in line with the *Enterobacteriaceae* bloom observed in natural-like microcosm experiments (Fig. 2A, C). To examine whether the age-dependent bloom was specific to gut *Enterobacteriaceae* or also involved increases in the abundance of other members of the gut microbiome, we used colony forming unit (CFU) counts and quantitative (q)PCR to evaluate gut bacterial load. In worms aging on CeMBio, an increase in total bacterial load (CFUs on non-selective LB) was observed, from 10^3^ bacterial cells/worm at D0 (L4 larvae) to around 10^5^ cells/worm at D12 of adulthood. However, a steeper increase was observed in the *Enterobacteriaceae* load (CFUs on selective VRBG plates) (Fig. 2B), from being barely detectable in D0 to about 5% of the total gut microbiome by old age (∼5000 cells/worm). A similar trend was observed in worms raised on the *Enterobacteriaceae-*rich SC20 community, as demonstrated with qPCR using universal *Eubacteria* primers or *Enterobacteriaceae-*specific primers and normalized to worm DNA (represented by actin genes). This evaluation showed a much steeper increase in *Enterobacteriaceae* strains compared to the increase in the total bacterial load (Fig. 2D). These results support the notion that an *Enterobacteriaceae* bloom is a hallmark of aging, regardless of the initial conditions such as high or low environmental diversity, or high or low initial proportion of *Enterobacteriaceae*. This bloom involves a large increase in total bacterial load, but a proportionally larger increase in the *Enterobacteriaceae* load per worm.

**Table 1:** SC20 members.

SC20 members
| Strain name | Family | genus | species |
| --- | --- | --- | --- |
| CEent1-RFP | *Enterobacteriaceae* | *Enterobacter* | *hormaechei* |
| MSPm1 | *Pseudomonadaceae* | *Pseudomonas* | *Berkeleyensis* |
| Cre-5.2 | *Enterobacteriaceae* | *Enterobacter* | NA |
| L3-3L | *Sphingobacteriaceae* | *Sphingobacterium* | *puteale* |
| oak-5.2 | *Enterobacteriaceae* | *Buttiauxella* | NA |
| CEent3-GFP | *Enterobacteriaceae* | *Enterobacter* | *ludwigii* |
| WG-2.2 | *Pseudomonadaceae* | *Pseudomonas* | *plecoglossicida* |
| WG-2.4 | *Micrococcaceae* | *Arthrobacter* | NA |
| WG-2.5 | *Microbacteriaceae* | *Microbacterium* | *barkeri* |
| 2.1 | *Bacillaceae* | *Bacillus* | *nealsonii* |
| 3.1 | *Bacillaceae* | *Bacillus* | *asahii* |
| 3.2 | *Bacillaceae* | *Bacillus* | *megaterium* |
| 10.1 | *Bacillaceae* | *Lysinibacillus* | *pakistanensis* |
| 10.3 | *Bacillaceae* | *Lysinibacillus* | *pakistanensis* |
| 14.1 | *Bacillaceae* | *Bacillus* | *subtilis* |
| 19.1.7 | *Bacillaceae* | *Lysinibacillus* | *fusiformis* |
| 19.3.3 | *Enterobacteriaceae* | *Rahnella* | *victoriana* |
| 19.3.8 | *Enterobacteriaceae* | *Buttiauxella* | *agrestis* |
| CBent2 | *Enterobacteriaceae* | *Lelliottia* | *amnigena* |
| Cbent1 | *Enterobacteriaceae* | *Citrobacter* | *freundii* |

**Figure 2:**
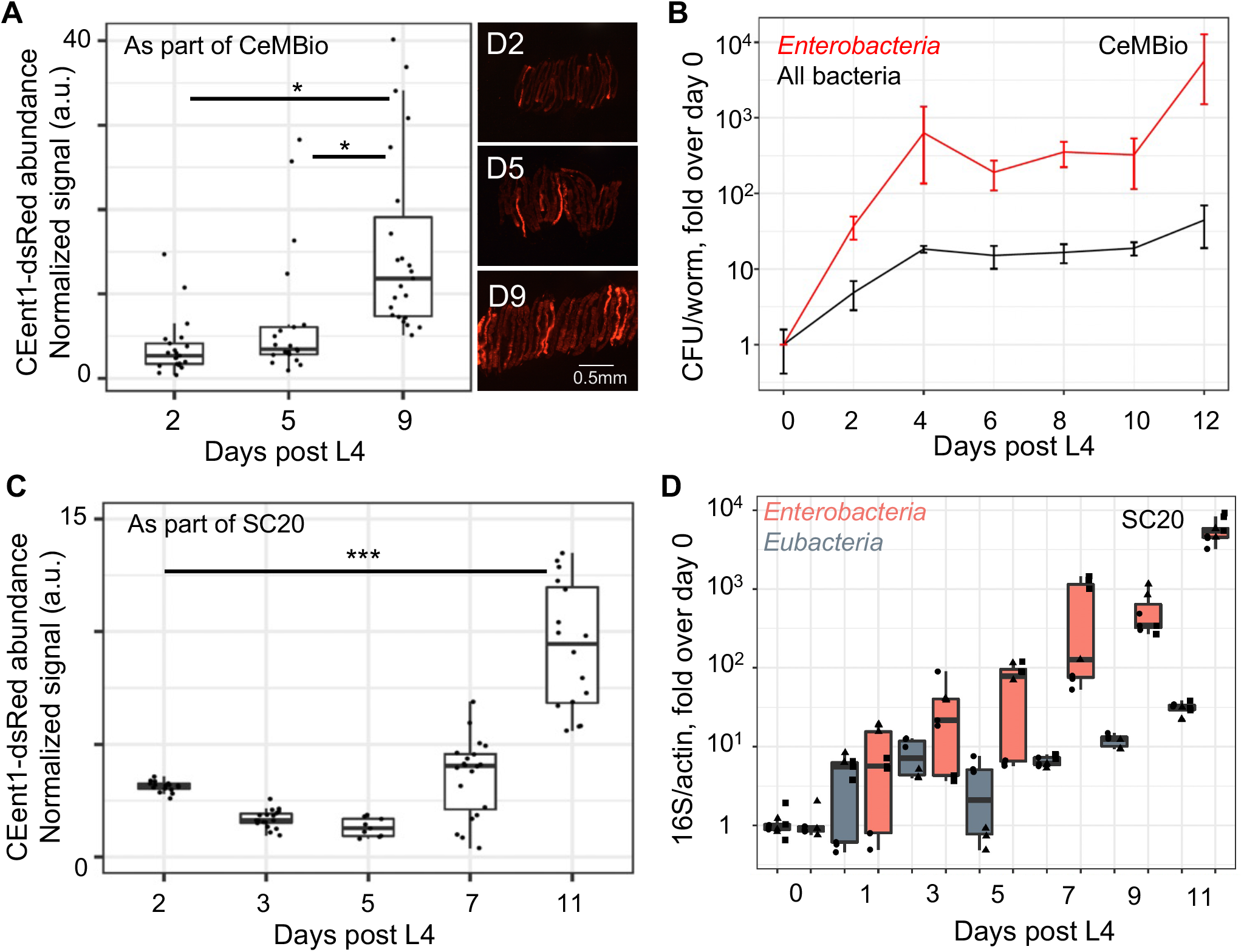
An *Enterobacteriaceae* bloom is a common denominator of aging worms raised in different microbial environments. **A**. Colonization of individual aging worms raised on CeMBio with CEent1*dsRed*, n=21-23/group; p < 0.01, pairwise t-tests; **B**. Bacterial load in aging worms raised on CeMBio, based on CFU counts of *Enterobacteriaceae* on VRBG plates and of total bacteria on LB plates. Shown are averages ± SDs for 3 plates per time point (n=4-12 worms/time point). **C**. CEent1 colonization in aging worms raised on SC20 with CEent1*dsRed* (n=9-18 worms*/*time point); p <0.001, pairwise t-tests. **D**. Fold change in bacterial load in worms aging on SC20, assessing bacterial load with qPCR using primers specific for *Enterobacteriaceae* 16S or Eubacterial 16s, normalized to worm DNA assessed by qPCR with primers specific for *C. elegans* actin (shapes represent replicate plates, each evaluated by qPCR in duplicate or triplicate).

### An *Enterobacteriaceae* bloom in aging animals can have detrimental consequences

Previous work showed that CEent1, serving as a representative of gut *Enterobacteriaceae*, protected young animals from infection with the pathogen *Enterococcus faecalis* ^22,30^. However, in worms disrupted for DBL-1/BMP immune signaling, gut abundance of CEent1 increased and the otherwise beneficial commensal became an exacerbating factor in host health outcomes ^25^. We thus used CEent1 to examine the functional significance of the age-dependent *Enterobacteriaceae* bloom. Worms raised on CEent1 and shifted to *E. faecalis* at the end of larval development showed higher pathogen resistance compared to worms raised on the *E. coli* control, as previously shown. In contrast, worms raised on CEent1 to middle age before shifting to *E. faecalis* (day four of adulthood, at which stage worms are well colonized), showed significantly lower pathogen resistance compared to worms raised on *E. coli* controls (Fig. 3A, B). Thus, the *Enterobacteriaceae* bloom has the potential to have detrimental consequences in aging worms.

**Figure 3:**
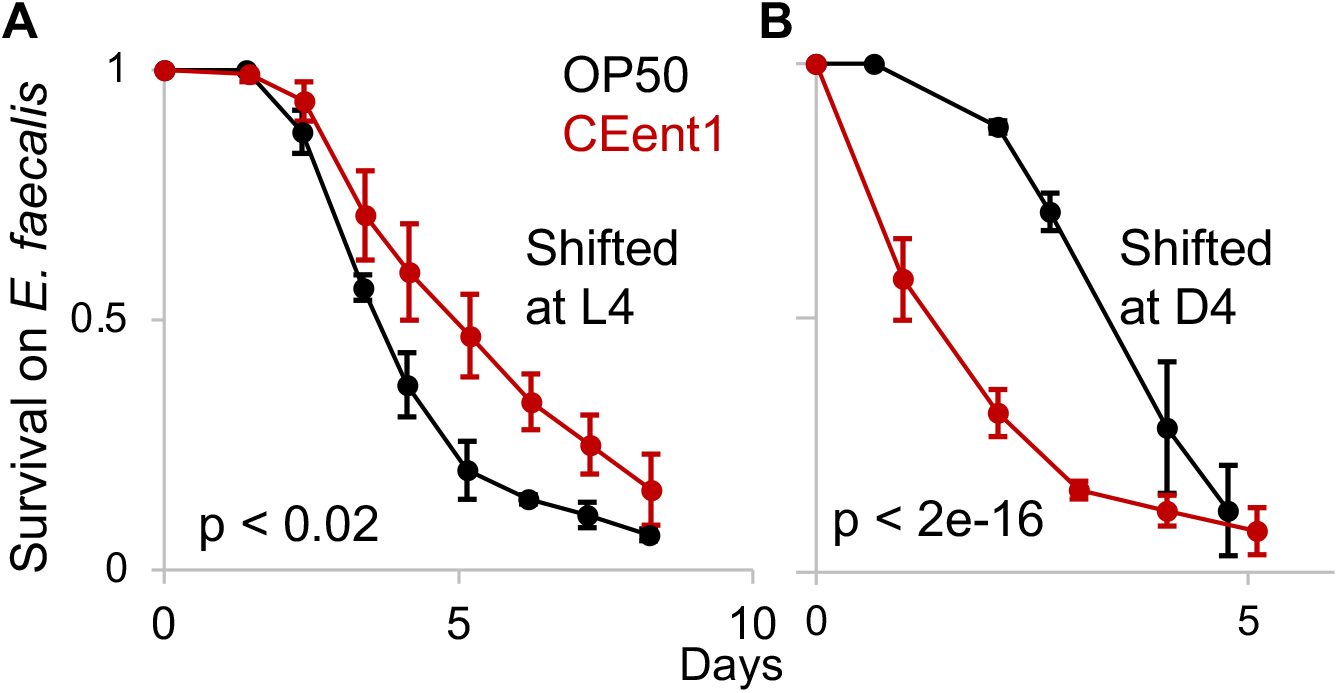
*Enterobacter hormachei* CEent1 bloom in aging worms is associated with increased susceptibility to infection. Survival curves for wildtype worms raised on designated monocultures and shifted to to plates with the pathogen, *E. faecalis* at L4 (**A**, n = 85-99/group), or at day four of adulthood (**B**, n = 96-99). p-values calculated with logrank test.

### Changes in the intestinal niche associated with an age-dependent decline in DBL-1/BMP signaling may underlie the *Enterobacteriaceae* bloom

What may be the cause for the *Enterobacteriaceae* bloom? Experiments in microcosm environments and with defined communities showed that environmental availability was not a likely cause (Fig. 1B, 2A-D). Ecological succession, driven by accumulating effects of interactions over time, also did not appear to contribute to the expansion (Fig. 1C). This was further supported by a comparison of CEent1-dsRed colonization in worms raised continuously on CEent1-dsRed monocultures versus worms shifted to CEent1-dsRed in different ages for a fixed duration of two days, which showed comparable age-dependent increases in *Enterobacteriaceae* abundance (Fig. 4A). We further examined whether age-dependent decline in bacterial uptake may play a role in causing the bloom. To this end, we compared CEent1-dsRed colonization, as part of the SC20 community, in wildtype worms, which show an age-dependent decline in pharyngeal pumping (and thus bacterial uptake), and *eat-2* mutants, which lack a pharyngeal receptor for the neurotransmitter acetylcholine resulting in slow pumping rate, which does not change considerably during aging (Fig. 4B inset). In both strains we observed a similar course of age-dependent increases in CEent1-dsRed colonization (Figure 4B), indicating that age-dependent changes in bacterial uptake likely did not contribute to the *Enterobacteriaceae* bloom.

**Figure 4:**
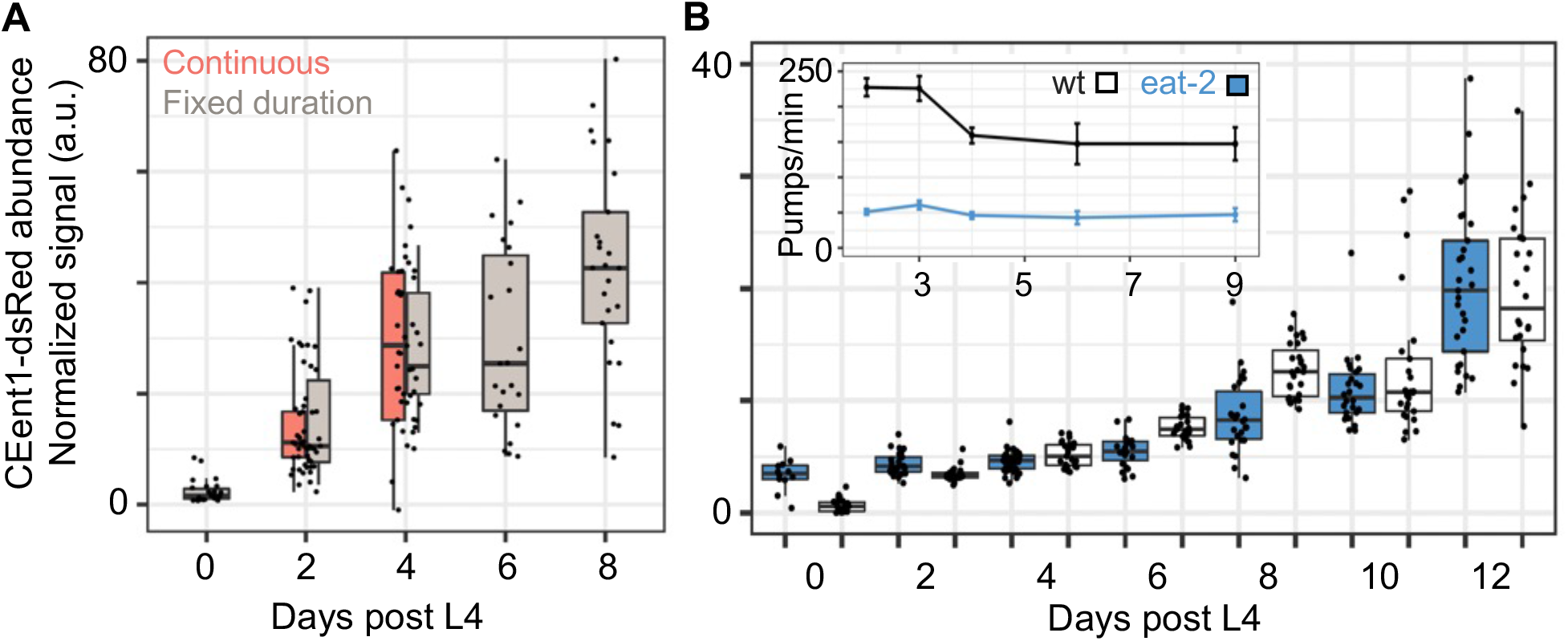
Neither duration of exposure to bacteria nor their rate of uptake contributes to the *Enterobacteriaceae* bloom. **A**. Colonization of aging worms by CEent1-dsRed following continuous exposure from larval stages or shifting worms for two days prior to the designated time points (fixed duration) (n=22-27 worms/group/time point). Box and whisker plots show median values, marked with a line, 25^th^ and 75^th^ percentile values delineating the box. **B**. CEent1*dsRed* colonization in wildtype and *eat-2* worms raised on the SC20 community (n = 10-36 worms/group/time point); **inset** demonstrates age-dependent declines in pumping rates; n = 4-10 worms/time point).

DBL-1/BMP signaling is a conserved regulator of development, body size and immunity ^31^. Previous work at the lab identified a role for DBL-1-dependent immune regulation in shaping the worm gut microbiome, particularly affecting *Enterobacter* strains, which bloomed when genes encoding different components of this pathway were disrupted ^25^. Gene expression data in Wormbase (https://wormbase.org) suggested that expression of the pathway’s components may decline at the end of larval development. To examine whether this indicated an age-dependent decline in DBL-1 signaling and downstream gene expression, which might affect the gut microbiome, we used a transgenic worm strain expressing GFP from the *spp-9* promoter, previously shown to be negatively regulated by DBL-1 signaling ^32^. Fluorescent imaging demonstrated age-dependent increase in the expression of the GFP reporter, indicating a decline in DBL-1 signaling in aging worms (Fig. 5A). In line with this, the effects of either disruption or over-expression of the *dbl-1* ligand gene diminished with age. Reduced effects of *dbl-1* disruption were also observed in worm colonization with CEent1, which in middle-aged mutants was comparable to that seen in wildtype animals, indicating a decline in DBL-1 signaling and in its involvement in controlling gut *Enterobacteriaceae* abundance during early aging (Fig. 5B). Further support for a decline in DBL-1 control of gut bacteria was provided by experiments with *sma-4(syb2546)* mutants, which carry a gain-of-function (gof) mutation in DBL-1’s transcriptional mediator, exhibiting a 20% longer body length compared to wildtype animals (Fig. 5C inset). These mutants showed lower CEent1-dsRed colonization compared to wildtype animals in early days of adulthood, up to day five (blue box compared to dotted line in Fig. 5C), signifying a delay in the CEent1 bloom. The ability to delay the CEent1 bloom had beneficial consequences, as sma-4(gof) mutants were partially protected from CEent1’s detrimental effects on infection resistance in day four adults (Fig. 5D). Together, these results suggest that an age-dependent decline in DBL-1 signaling alters the intestinal niche, permitting preferential accumulation of *Enterobacteriaceae*, which can be detrimental. Boosting DBL-1 signaling may mitigate the bloom and its consequences, but only partially.

**Figure 5:**
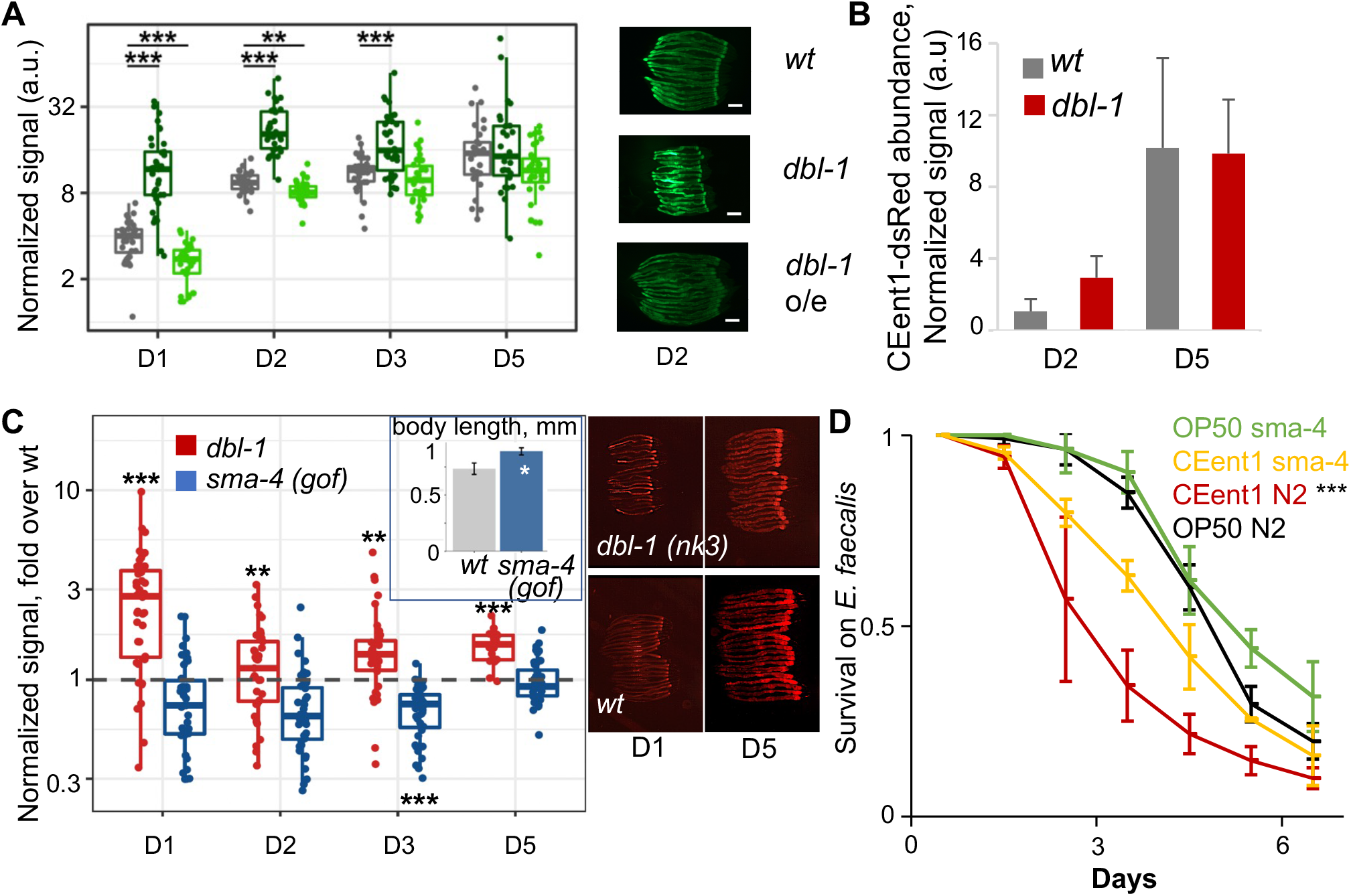
The *Enterobacteriacae* bloom is associated with an age-dependent decline in DBL-1/BMP signaling. **A**. GFP expression from the *spp-9* promoter in designated strains; representative images (scale bar = 200 µM) and quantification (n = 27-37/group/time point); \*\**p* < 0.01, and \*\*\**p* < 0.001, Kruskal-Wallis rank sum test and *post hoc* Wilcoxon test (mutant *vs* wt), Bonferroni corrected. **B**. CEent1*dsRed* colonization in worms of designated strains at designated days of adulthood. **C**. *dbl-1* null mutants, and *sma-4* gof mutants aging on CeMBio containing CEent1-dsRed (n = 39-40/group/time point); fold over wt median. **Inset**. Body length of designated strains (n=9-12, p < 0.0001) **D**. Survival of worms of designated strains raised on *E. coli* or *CEent1* and shifted to *E. faecalis* at D4 of adulthood (n = 106-112/group); ***, p < 0.001, logrank test.

### Commensal communities can effectively mitigate the detrimental consequences of the *Enterobacteriaceae* bloom

The ability of the *sma-4(gof)* mutation to partially mitigate the detrimental effects of CEent1 expansion suggested that protection from an *Enterobacteriaceae* bloom is possible. Considering that the decline in DBL-1 signaling also increased abundance of non-*Enterobacteriaceae* bacteria (Fig. 2), we examined whether other bacteria could compete with CEent1 and help prevent its detrimental effects. Recent work identified members of the genus *Pantoea* as common worm commensals effectively colonizing the gut and capable of competing with an invading pathogen ^23^. Wildtype worms raised on a community consisting of three such *Pantoea* commensals in addition to CEent1-dsRed (in equal parts) and shifted to *E. faecalis* in middle-age were as resistant as worms raised on *E. coli* alone, and significantly more resistant than worms raised on a similar inoculum of CEent1-dsRed mixed with *E. coli* (Fig. 6A). While mortality on *E. faecalis* plates was attributed to the pathogen, 96.7% of the worms raised on the CEent1-dsRed/*E. coli* mix, which died in any of the days of the infection assay, were heavily colonized with CEent1-dsRed (Fig. 6A inset), indicating proliferation alongside *E. faecalis*. In contrast, only 62.5% of the worms who were initially raised on the CEent1-dsRed/*Pantoea* mix were colonized, together indicating that the *Pantoea* community was able to mitigate CEent1 proliferation in some of the worms and to reduce mortality in the population. To examine whether mitigating the detrimental effects of CEent1 proliferation was unique to *Pantoea*, worms were raised on a subset of seven members of CeMBio (see Methods), with or without BIGb393 (one of the protective *Pantoea* strains, which is also a member of CeMBio) and with an excess of CEent1-dsRed (50% of total) and shifted at middle age to *E. faecalis*. Raising worms on the CeMBio subsets, with or without BIGb0303, conferred significantly higher resistance to infection than in worms raised on CEent1-dsRed alone (Fig. 6B). Again, fewer of the dead worms were colonized with CEent1-dsRed among those raised on CeMBio/CEent1-dsRed (51.4%), compared to those raised on CEent1-dsRed alone (84.6%). These experiments demonstrate that over proliferation of *Enterobacteriaceae* and its detrimental consequences in aging worms can be mitigated with more than one combination of gut commensals. Lastly, in the context of such a community, CEent1, or other *Enterobacteriaceae*, did not compromise the lifespan of their host, as worms grown on CeMBio with or without its *Enterobacteriaceae* members (CEent1 and JUb66) had a comparable lifespan (Fig. 6C).

**Figure 6:**
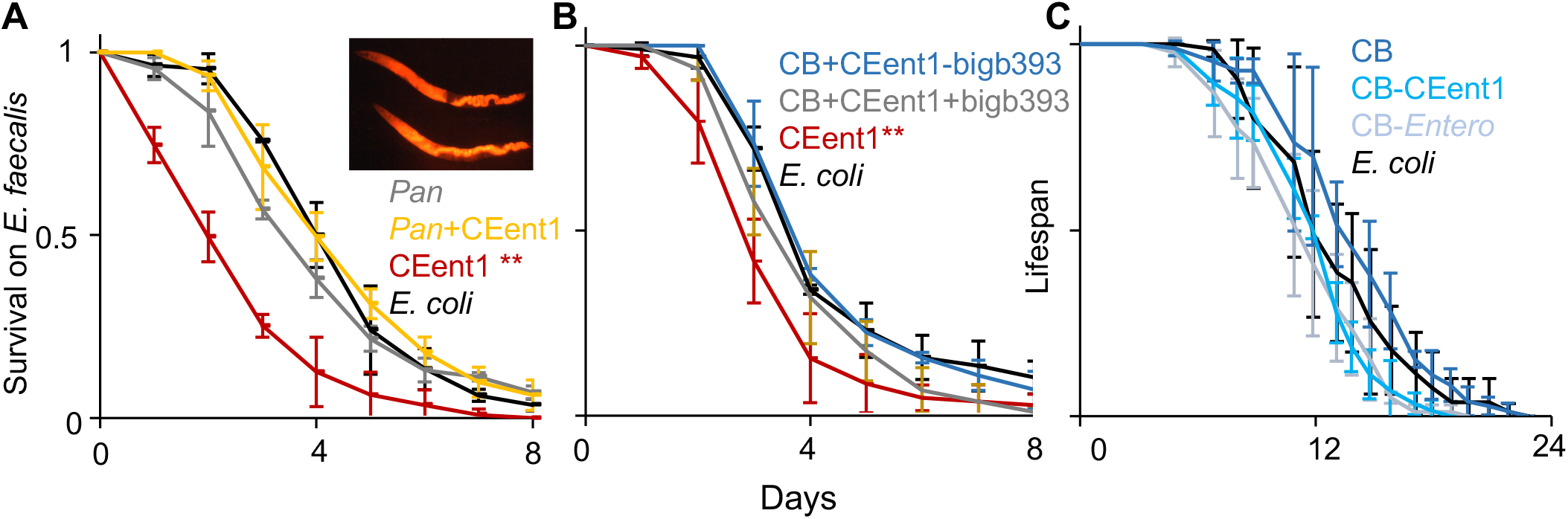
Commensal communities mitigate age-dependent susceptibility to infection. **A**. Survival of worms raised on the designated strains/communities and shifted to *E. faecalis* at D4; *Pan*, a community of three *Pantoea* strains; **, p < 0.0001, log rank test (n = 82-90/group); averages ± SDs for three plate replicates. **Inset**. CEent1-colonized dead worms, one day after shift to *E. faecalis*. **B**. BIGb*393*, a *Pantoea* strain in CeMBio.(CB) **, p < 0.0001, n = 95-105/group). Shown are results of one representative experiment out of two with similar results. **C**. Lifespan of wildtype worms raised on designated communities. *Entero* stands for *Lelliotia* Jub66 and *Enterobacter* CEent1. Averages ± SDs for three plate replicates (n = 67-79/group).

## Discussion

Our experiments identify an *Enterobacteriaceae* bloom as a hallmark of the gut microbiome in aging *C. elegans*. This bloom was observed in worms raised in natural-like microcosm environments with varying initial microbial diversity, as well as in worms raised on defined bacterial communities differing in the environmental availability of *Enterobacteriaceae*, indicating that it is independent of initial conditions. The *Enterobacteriaceae* bloom is not due to bacteria-driven ecological succession or to age-dependent changes in bacterial uptake. Rather, it is due to intrinsic age-dependent changes in the intestinal niche, suggesting that the bloom is a signature of chronological age. Our results demonstrate that increased gut abundance of *Enterobacteriaceae* strains may have detrimental consequences for aging animals, at least for infection resistance. However, in the context of a community, even such with restricted diversity, the detrimental consequences of this bloom can be mitigated. Our results highlight the *Enterobacteriaceae* bloom as a hallmark of chronological aging but suggest that the consequences of this bloom are context-dependent, with microbiome composition representing the context that can differentiate between healthy or unhealthy aging.

It is accepted that aging is accompanied by gut dysbiosis ^7–9,14,33^. Human studies have documented diverse changes in gut microbiome composition. Among those, increased abundance of *Proteobacteria/Pseudomonadota*, and specifically of *Enterobacteriaceae*, is a recurring theme ^17,34^. In line with this, our results show a replicable age-associated expansion of *Enterobacteriaceae*, suggesting that it may be an evolutionarily conserved signature of aging. What causes this bloom is not clear. A study in fruit flies describes gut dysbiosis characterized by a biphasic change in the microbiome of aging flies, in which a midlife bloom of *γ-proteobacteria* led to intestinal barrier dysfunction, and a subsequent increase in *α-proteobacteria* ^7^. Whereas barrier dysfunction was suggested as the cause for gut dysbiosis ^35^, what initiated the *γ-proteobacteria* bloom was not clear. The *Enterobacteriaceae* bloom we observed in worms may be analogous to the initial phase of dysbiosis in flies. In worms, initiation of the *Enterobacteriaceae* bloom was associated with a decline in DBL-1/BMP signaling. DBL-1 signaling was previously shown to play a central role in controlling gut *Enterobacteriaceae* ^25^. Thus, its age-dependent decline may change the host intestinal niche making it more permissible for *Enterobacteriaceae* expansion. Supporting a causal role for DBL-1 signaling in the *Enterobacteriaceae* bloom, gain-of-function mutants for the SMA-4 BMP mediator showed a delay in *Enterobacter* colonization as well as attenuated infection susceptibility. These results suggest that decline of immune signaling during aging is an important factor in initiating dysbiosis. However, at least in the case of SMA-4, revamping the immune pathway to mitigate the detrimental effects of the bloom was only partially successful, suggesting that additional changes in the gut niche may take place to promote the *Enterobacteriaceae* bloom, and further limiting the potential of DBL-1/BMP reactivation in aging worms as an intervention to alleviate gut dysbiosis.

*Enterobacteriaceae* blooms are associated with increased susceptibility to infection ^36–39^. In agreement with this, expansion of *E. hormachei* CEent1 in aging worms compromised infection resistance and survival. Continued gut accumulation of CEent1 following a shift to pathogen plates further suggests that in old worms CEent1 (and perhaps additional *Enterobacteriaceae* members) was an opportunistic pathogen, or a pathobiont. Previous results support the notion that CEent1 was a pathobiont, as worms growing on CEent1 alone have a shorter lifespan compared to a those raised on an *E. coli* diet ^22^. However, in the context of a community (CeMBio), inclusion or removal of CEent1 did not have any effect on lifespan, supporting the importance of a diverse microbiome in keeping pathobionts in check as the host ages.

The genetic tractability and short lifespan of *C. elegans* has made it a useful model for aging research. Its more recent establishment as a model for microbiome research adds to that the advantages of longitudinal microbiome analysis in clonal host populations, while having greater control over bacterial availability, to facilitate studies of host-microbiome interactions during aging. Using this model, we identified what seems to be an evolutionary conserved signature of dysbiosis in aging animals and have begun to dissect its causes as well as its consequences. As often seen in different scenarios of gut dysbiosis, the *Enterobacteriaceae* bloom that we identified is associated with pathology. However, this pathology can be circumvented by manipulating the gut microbiome using various commensal communities. Thus, while an *Enterobacteriaceae* bloom seems to be an inevitable consequence of aging, its extent and outcomes can be restrained by other members of the gut microbiome. What differentiates between communities that can or cannot achieve this remains to be seen.

## Materials and methods

### Worm strains

*C. elegans* strains included wildtype N2, *dbl-1(nk3)*, and the *dbl-1* overexpressing strain BW1940, obtained from the *Caenorhabditis* Genome Center (CGC); transgenic strains expressing GFP from the *spp-9* promoter: TLG690, texIs127[spp-9p::GFP], TLG707, texIs127;*dbl-1*(*nk3*) and TLG708, texIs127;texIs100[*dbl-1*p::GFP::*dbl-1*], were gratefully received from Tina Gumienny ^32,40^; gain of function (gof) *sma-4(syb2546)* mutants were gratefully received from Cathy Savage-Dunn; and *eat-2(ad1116)* mutants were gratefully received from Andrew Dillin.

### Bacterial strains and communities

*Escherichia coli* strain OP50 and the Gram-positive pathogen, *Enterococcus faecalis* strain V583 were obtained from the CGC. Two defined communities of worm gut commensals were used: CeMBio, with twelve strains, represents the worm core gut microbiome ^29^ and SC20, a subset of twenty strains of the previously described SC1 ^25^, with eight species out of the 20 of the *Enterobacteriaceae* family (Table 1). In addition, a subset of of CeMBio strains was used in experiments testing effects on susceptibility to *Enterococcus faecalis* infection, including only strains that are sensitive to gentamycin, which is used in *E. faecalis* plates, to prevent enrichment of gut commensals through the environment. Additional commensals, of the genus *Pantoea*, family *Erwiniaceae* (a recent splinter off *Enterobacteriaceae)*, included BIGB0393 (also in CeMBio) and the recently characterized *Pantoea cypripedii* strains V8 and T16 ^23^.

Bacterial communities were prepared for experiments by growing individual strains in LB at 28°C for two days, adjusting cultures to 1 OD, concentrating 10-fold and mixing equal volumes from each culture. 100-200 µL aliquots of the mix were plated on either minimal nematode growth medium (NGM) or on peptone-free medium (PFM), which further limits bacterial growth ^29^, as described, and air-dried for 2 to 12 hours prior to the addition of worms.

### Construction of fluorescently-tagged *Enterobacter hormaechei* CEent1-dsRed

*E. hormaechei* CEent1, previously misidentified as *E. cloacae* ^22^, is a member in both the CeMBio and SC20 communities. The construction of its dsRed-expressing derivative was achieved by integrating the *dsRed* gene into the functionally neutral *attTn7* site in the CEent1 genome using the site-specific *Tn7-*mini transposon system ^41^ (Supplementary Fig. 1). Transposon insertion was achieved through triparental mating with a donor strain and a transposase-expressing helper strain, both based on the DAP-auxotrophic and *pir1*-positive *E. coli* strain BW29427, containing the conjugative RP4 mating system as a chromosomal insert. This strain, as well as other *E. coli* strains and plasmids used in constructing the fluorescently-tagged strains, were gratefully received from the Goodrich-Blair lab, University of Tennessee Knoxville. Briefly, the original *GFP* in the mini-transposon, carried on plasmid pURR25 (Supplementary Fig. 1), was replaced with *dsRed* by digesting the plasmid with BseRI and NheI to cleave out the *GFP* coding sequence ^42^, amplifying the *dsRed* gene from pBK-miniTn7-ΩGm-DsRed ^43^ using primers 5’-TAC GTG CAA GCA GAT TAC GG-3’ and 5’-ATC CAG TGA TTT TTT TCT CCAT-3,’ and ligating the amplified *dsRed* to the linearized pURR25 vector. The modified pURR25 plasmid, carrying the pir1-dependent *oriR6K*, as well as *dsRed* and the antibiotic resistance genes *KanR* and *StrR*, was re-introduced by electroporation into its original host strain, constituting the donor strain. The transposase plasmid, pUX-BF13, in the helper strain, also included the pir1-dependent *oriR6K* and *AmpR* antibiotic resistance, along with the *Tn7-*transposase ^44^.

Recipient strain CEent1 (*pir1*-negative), donor, and helper strains were each cultured until they reached an OD_600_ of 0.4 and then mixed in a 1:1:1 ratio in SOC DAP media for one hour at 37 °C. The mixture was spread on LB DAP plates for an additional 24-hour incubation to promote conjugation. Bacteria subsequently underwent several rounds of re-streaking on Kan^+^/DAP^−^ plates to select for integrant CEent1 cells and to dilute-out the *pir1-*negative plasmids which cannot be replicated in CEent1 (Supplementary Fig. 1). Integrant clones were verified as CEent1 by sequencing a 200bp fragment of the CEent1 gene for *gyrB* using the primers 5’-GCA AGC AGG AAC AGT ACA TT-3’ and 5’-TCG GCT GAT AAA TCA GCT CTT TC-3’.

### Microcosm experiments

Compost microcosms harboring diverse microbial communities were prepared from local soil composted with produce for up to two weeks essentially as previously described ^19,45^. Briefly, local soils were supplemented with banana peels or chopped apple, composted soils were split into two parts: one part (6 gr in a glass vial) autoclaved to eliminate native nematodes and the other (10 gr soil) suspended in M9 buffer to obtain a microbial extract which was concentrated and added to the autoclaved samples to reconstitute the original microbial community.

In continuous aging experiments, synchronized populations of germ-free L1 worms were raised at 20°C in separate vials containing the same compost and harvested at advancing ages up to day five of adulthood (D5) (Fig. 1A). The final time point was determined by the need to distinguish between the original cohort (post-gravid at D5) and progeny (mid-stage gravids), which could not be achieved in subsequent time points. In experiments with fixed time colonization, worms were raised on live *E. coli* until the L4 stage to ensure proper development, then transferred to kanamycin-killed *E. coli* ^46^ from which worms were further transferred at advancing ages to microcosm environments for two days before harvesting for analysis. For the earliest time point (gravids, day zero of adulthood), worms were raised on live *E. coli* from L1 to the L4, then shifted to dead *E. coli* for 4 hours to minimize carry over of live *E. coli*, before transferring to microcosms. Worm harvesting was carried out using a Baermann funnel as described ^45^. Soil samples (1g) were taken from microcosms of the same compost batch used to grow worms (“soil”), or from the same microcosm from which worms were harvested (“grazed soil”).

### Experiments with defined bacterial communities

Aging experiments on bacterial communities/strains were carried out similarly to the description for microcosm experiments, with bacteria seeded on NGM or PFM plates as described. In continuous aging experiments, worms were transferred to fresh plates every day during the reproductive phase to separate the original cohort from their progeny, enabling carrying on experiments into later stages of adulthood.

### DNA extraction

Worms harvested in microcosm experiments (100-500 per group) were extensively washed, surface-sterilized by letting them crawl for an hour on plates with 100 µg/mL gentamycin and used for DNA extraction with the QIAGEN PowerSoil DNA isolation kit (Cat. #12888) as previously described ^25^.

Worms harvested from plates with defined communities (100-150 per group) were washed three times with M9, paralyzed with 25mM levamisole (Acros Organics) to close their intestine, and surface-sterilized with 2% bleach in M9, as described ^45^. DNA was extracted using the DNeasy PowerSoil Pro Kit (Qiagen Cat. # 47016) according to manufacturer instructions, with the following modification: to break open worms prior to the first step of the protocol, worms were incubated in the kit’s buffer for 10 minutes at 60°C, then crushed with added zirconium beads using a PowerLyzer (2000 RPM 2 × 30 seconds).

### 16S Next Generation Sequencing (NGS) and analysis

DNA samples from worms and their respective environments were used to generate sequencing libraries of the 16S V4 region. Libraries from microcosms experiments were prepared using tailed primers, as previously described ^19^, and sent for 150 paired-end sequencing to the UC Davis sequencing facility. Analysis of bacterial 16S amplicon data was carried out using the QIIME2 software pipeline ^47^. Sequence reads were demultiplexed and filtered for quality control, with 85.6% of all reads passing quality filtering, providing an average read of 121,053 reads per sample. Sequences were aligned and clustered into operational taxonomic units (OTU) based on the closed reference OTU picking algorithm using the QIIME2 implementation of UCLUST ^48^, and the taxonomy of each OTU was assigned based on 99% similarity to reference sequences based on Greengenes release 13_8. Prior to diversity analysis, all communities were rarefied to 116,543 sequences per sample. Shannon’s diversity and Faith’ phylogenetic diversity were calculated to assess *alpha* diversity of soil and worms gut microbiotas ^49,50^. Shannon diversity is a composite measure of both richness and evenness, while Faith method takes the phylogenetic distance of species into account. Weighted UniFrac distances were calculated to assess *beta* diversity and used in principal coordinates analysis (PCoA). Raw data and metadata can be accessed at http://www.ncbi.nlm.nih.gov/sra with accession no.: PRJNA982115.

### Fluorescence imaging

For each time point examined, 30-40 worms were picked off CEent1-dsRed, or CEent3-GFP containing plates (with the single strain, or with the fluorescent strains as part of a community), washed with M9, paralyzed with 10 mM Levamisole, and mounted on a slide with a 2-4% agarose pad. Imaging was performed with a Leica MZ16F equipped with a QImaging MicroPublisher 5.0 camera. Quantification of fluorescent signal was conducted using the Fiji plugin within the ImageJ package v2.10/1.53c or 1.53f51 ^51^. Integrated Density values were measured per worm after subtracting background mean gray values and autofluorescence and normalized for worm area.

### Quantifying bacterial load by colony forming unit (CFU) counting

For each time point examined 10-15 worms were washed three times in M9-T (M9/0.0125% TritonX-100), paralyzed with 25 mM Levamisole, and surface sterilized by a three-minute incubation in 2% bleach ^29^. After two washes of 1mL of PBS-T (PBS/0.0125% Triton-X 100), worms were collected in a final volume of 100µL. Samples of final wash were plated and incubated at 28°C for 2 days to verify removal of external bacteria. To release bacteria from the worm gut, worms were crushed with 10-15 zirconium beads in 100µL of PBS in a PowerLyzer at 4000 RPM for 30 to 45 seconds. Released bacteria were diluted and plated either on non-selective LB agar plates (for all bacteria) or *Enterobacteriaceae-*selective VRBG plates. CFUs were counted after 1-2 days of incubation at 28°C. CFUs for *E. hormaechei* and *Lelliottia amnigena* could be distinguished on VRBG plates based on morphology.

### Quantifying bacterial load via qPCR

Relative bacterial load was measured with real-time quantitative PCR using the eubacterial 16S rDNA primers 806f (5’-AGATACCCCGGTAGTCC-3’) and 895r (5’-CYGYACTCCCCAGGYG-3’), and the *Enterobacteriaceae-*specific 16S specific primers Ent_MB_F (5’-ACCTGAGCGTCAGTCTTTGTC-3’) and R (5’-GTAGCGGTGAAATGCGTAGAGA-3’) ^25^. qPCR was performed using Bio-Rad SsoAdvanced Universal SYBR Green qPCR Supermix and an Applied Biosystems StepOne Plus real-time PCR system. Cycling (eubacterial primers): 95°C for 5 min, 45 x [95°C for 15 sec, 60°C for 30 sec, 72°C for 15 sec], 72°C for 5 min; and for the *Enterobacteriaceae* specific primers: 95°C for 5 minutes, 40 x [95°C for 15 sec, 60°C for 30 sec], 72°C for 5 minutes. Ct values for bacterial 16S were normalized to worm material by subtracting Ct values obtained for worm actin using the pan-actin primers ^52,53^.

### Survival assays

For infection resistance experiments, synchronized worm populations were raised from L1 on PFM plates with *E. coli* OP50, CEent1, or designated communities, and shifted, at L4, or at day 4 of adulthood, to *E. faecalis* plates prepared with Brain Heart Infusion Agar containing 25 µg/µL gentamicin and seeded with bacteria a day before the transfer of worms. Assays were carried out at 25°C and dead or live worms were counted every day ^30^. For lifespan assays, worms were raised, on designated strains or communities in PFM plates at 20°C and scored daily for survival beginning at L4 (t_0_).

### Statistical analyses

Statistical tests were conducted in R (v 3.6.3). Survival curves were statistically compared using Kaplan-Meier analysis and log-rank tests using the survdiff R package ^54^ and all graphs were created with the ggplot R package ^55^.

## Acknowledgments

We thank Dr. Heidi Goodrich-Blaire for providing plasmids required for constructing the dsRed-expressing *Enterobacter hormaechei*, Dr. Tina Gumienny for *spp-9* reporter strains, to Dr. Cathy Savage-Dunn for *sma-4* mutants, and to Dr. Andrew Dillin for *eat-2* mutants.

Work described in this manuscript was supported by NIH grants R01OD024780 and R01AG061302. D.K was supported by NSF fellowship DGE 2146752; J.C. was supported by a fellowship from Berkeley’s Center for Research in Aging.

## Contributions

R.C. M.B. and M.S. conceived the project, and M.S. supervised it; M.B. conducted microcosm experiments which were analyzed by D.K. R.C., R.B., B.P., D.M., E.D. and J.C. carried out experiments and analyses. V.N. generated the CEent1-dsRed strain. R.C. and M.S. compiled the results and wrote the manuscript.

## Ethics declarations

The authors declare that the research was conducted in the absence of any commercial or financial relationships that could be construed as a potential conflict of interest.

---

**Supplementary Figure 1:**
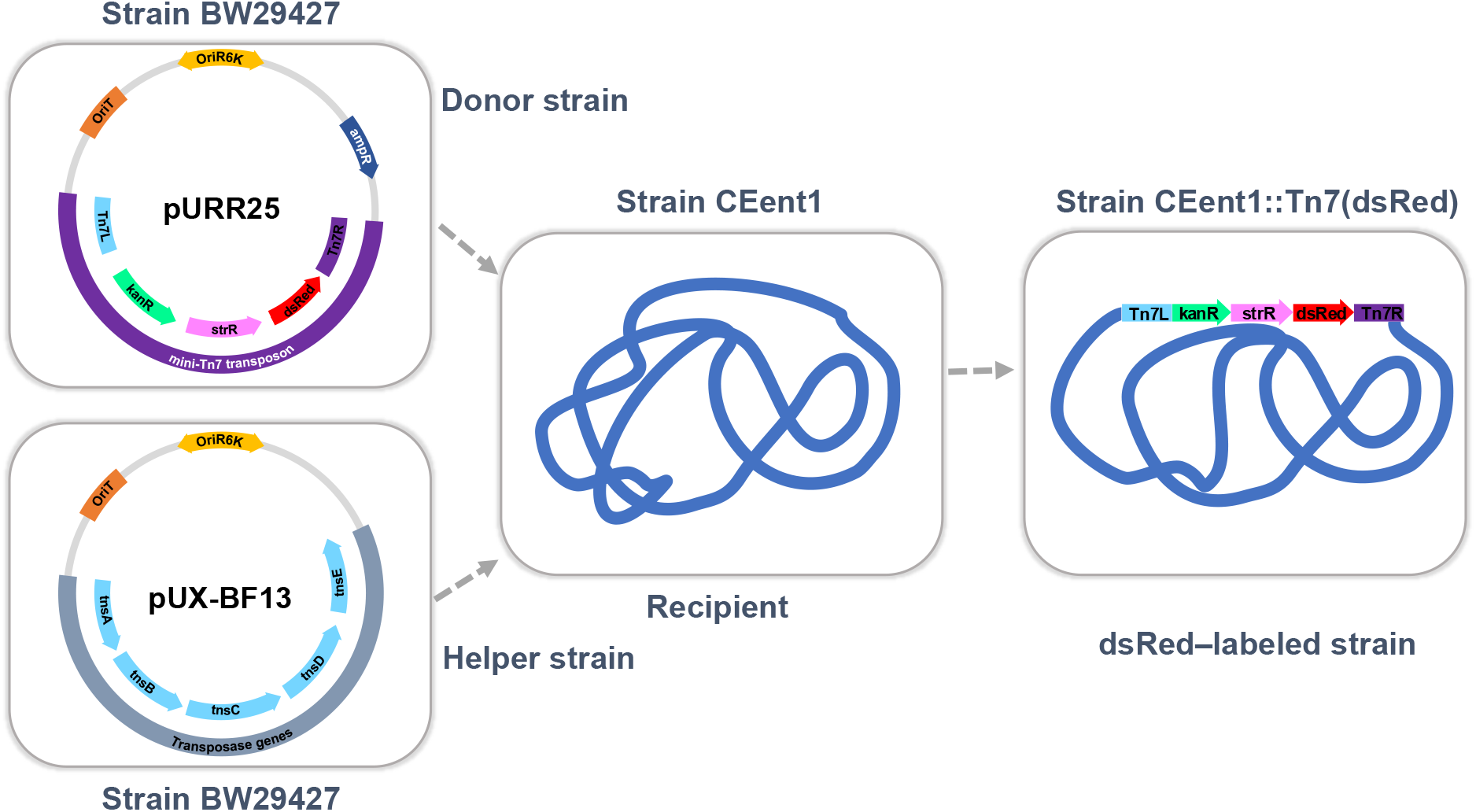
Plasmids used in triparental mating to generate dsRed-expressing *Enterobacter hormaechei* CEent1. Shown are maps of the *Tn7-gfp* donor plasmid (pURR25) and the transposase-carrying helper plasmid (pUX-BF13). Tn7R and Tn7L are the sites recognized and cut by the transposase.

## Notes

### Competing Interest Statement

The authors have declared no competing interest.

## References

1. Nicholson, J. K. et al. Host-Gut Microbiota Metabolic Interactions. Science 336, 1262–1267 (2012).

2. Dominguez-Bello, M. G., Godoy-Vitorino, F., Knight, R. & Blaser, M. J. Role of the microbiome in human development. Gut 68, 1108–1114 (2019).

3. Fan, Y. & Pedersen, O. Gut microbiota in human metabolic health and disease. Nat. Rev. Microbiol. 19, 55–71 (2021).

4. Morais, L. H., Schreiber, H. L. & Mazmanian, S. K. The gut microbiota–brain axis in behaviour and brain disorders. Nat. Rev. Microbiol. 19, 241–255 (2021).

5. Turnbaugh, P. J., Bäckhed, F., Fulton, L. & Gordon, J. I. Diet-Induced Obesity Is Linked to Marked but Reversible Alterations in the Mouse Distal Gut Microbiome. Cell Host Microbe 3, 213–223 (2008).

6. Mariño, E. et al. Gut microbial metabolites limit the frequency of autoimmune T cells and protect against type 1 diabetes. Nat. Immunol. 18, 552–562 (2017).

7. Clark, R. I. et al. Distinct Shifts in Microbiota Composition during Drosophila Aging Impair Intestinal Function and Drive Mortality. Cell Rep 12, 1656–1667 (2015).

8. O’Toole, P. W. & Jeffery, I. B. Gut microbiota and aging. Science 350, 1214–1216 (2015).

9. Thevaranjan, N. et al. Age-Associated Microbial Dysbiosis Promotes Intestinal Permeability, Systemic Inflammation, and Macrophage Dysfunction. Cell Host Microbe 21, 455-466.e4 (2017).

10. Wilmanski, T. et al. Gut microbiome pattern reflects healthy ageing and predicts survival in humans. Nat. Metab. 3, 274–286 (2021).

11. Biagi, E. et al. Gut Microbiota and Extreme Longevity. Curr. Biol. 26, 1480–1485 (2016).

12. Kim, K. H. et al. Gut microbiota of the young ameliorates physical fitness of the aged in mice. Microbiome 10, 238 (2022).

13. Parker, A. et al. Fecal microbiota transfer between young and aged mice reverses hallmarks of the aging gut, eye, and brain. Microbiome 10, 68 (2022).

14. Smith, P. et al. Regulation of life span by the gut microbiota in the short-lived African turquoise killifish. eLife 6, e27014 (2017).

15. Bárcena, C. et al. Healthspan and lifespan extension by fecal microbiota transplantation into progeroid mice. Nat. Med. 25, 1234–1242 (2019).

16. Adriansjach, J. et al. Age-Related Differences in the Gut Microbiome of Rhesus Macaques. J. Gerontol. Ser. A 75, 1293–1298 (2020).

17. Leite, G. et al. Age and the aging process significantly alter the small bowel microbiome. Cell Rep. 36, 109765 (2021).

18. Shapira, M. Host–microbiota interactions in Caenorhabditis elegans and their significance. Curr. Opin. Microbiol. 38, 142–147 (2017).

19. Berg, M. et al. Assembly of the Caenorhabditis elegans gut microbiota from diverse soil microbial environments. ISME J. 10, 1998–2009 (2016).

20. Dirksen, P. et al. The native microbiome of the nematode Caenorhabditis elegans: gateway to a new host-microbiome model. BMC Biol. 14, 38 (2016).

21. Zhang, F. et al. Caenorhabditis elegans as a Model for Microbiome Research. Front. Microbiol. 8, (2017).

22. Berg, M., Zhou, X. Y. & Shapira, M. Host-Specific Functional Significance of Caenorhabditis Gut Commensals. Front. Microbiol. 7, (2016).

23. Pérez-Carrascal, O. M. et al. Host Preference of Beneficial Commensals in a Microbially-Diverse Environment. Front. Cell. Infect. Microbiol. 12, (2022).

24. Slowinski, S. et al. Interactions with a Complex Microbiota Mediate a Trade-Off between the Host Development Rate and Heat Stress Resistance. Microorganisms 8, 1781 (2020).

25. Berg, M. et al. TGFβ/BMP immune signaling affects abundance and function of C. elegans gut commensals. Nat. Commun. 10, 1–12 (2019).

26. Taylor, M. & Vega, N. M. Host Immunity Alters Community Ecology and Stability of the Microbiome in a Caenorhabditis elegans Model. mSystems 6, e00608–20.

27. Zhang, F. et al. Natural genetic variation drives microbiome selection in the Caenorhabditis elegans gut. Curr. Biol. 31, 2603-2618.e9 (2021).

28. Langille, M. G. et al. Microbial shifts in the aging mouse gut. Microbiome 2, 50 (2014).

29. Dirksen, P. et al. CeMbio - The Caenorhabditis elegans Microbiome Resource. G3 GenesGenomesGenetics 10, 3025–3039 (2020).

30. Sifri, C. D. et al. Virulence Effect of Enterococcus faecalis Protease Genes and the Quorum-Sensing Locus fsr in Caenorhabditis elegans and Mice. Infect. Immun. 70, 5647–5650 (2002).

31. Savage-Dunn, C. & Padgett, R. W. The TGF-β Family in Caenorhabditis elegans. Cold Spring Harb. Perspect. Biol. 9, a022178 (2017).

32. Lakdawala, M. F. et al. Genetic interactions between the DBL-1/BMP-like pathway and dpy body size–associated genes in Caenorhabditis elegans. Mol. Biol. Cell 30, 3151–3160 (2019).

33. Claesson, M. J. et al. Composition, variability, and temporal stability of the intestinal microbiota of the elderly. Proc. Natl. Acad. Sci. U. S. A. 108, 4586–4591 (2011).

34. Odamaki, T. et al. Age-related changes in gut microbiota composition from newborn to centenarian: a cross-sectional study. BMC Microbiol. 16, 90 (2016).

35. Salazar, A. M. et al. Intestinal Snakeskin Limits Microbial Dysbiosis during Aging and Promotes Longevity. iScience 9, 229–243 (2018).

36. Lupp, C. et al. Host-Mediated Inflammation Disrupts the Intestinal Microbiota and Promotes the Overgrowth of Enterobacteriaceae. Cell Host Microbe 2, 119–129 (2007).

37. Bailey, M. T. et al. Stressor Exposure Disrupts Commensal Microbial Populations in the Intestines and Leads to Increased Colonization by Citrobacter rodentium. Infect. Immun. 78, 1509–1519 (2010).

38. Shaler, C. R. et al. Psychological stress impairs IL22-driven protective gut mucosal immunity against colonising pathobionts. Nat. Commun. 12, 6664 (2021).

39. Schlechte, J. et al. Dysbiosis of a microbiota–immune metasystem in critical illness is associated with nosocomial infections. Nat. Med. 29, 1017–1027 (2023).

40. Madhu, B., Lakdawala, M. F., Issac, N. G. & Gumienny, T. L. Caenorhabditis elegans saposin-like spp-9 is involved in specific innate immune responses. Genes Immun. 21, 301–310 (2020).

41. McKown, R. L., Waddell, C. S., Arciszewska, L. K. & Craig, N. L. Identification of a transposon Tn7-dependent DNA-binding activity that recognizes the ends of Tn7. Proc. Natl. Acad. Sci. 84, 7807–7811 (1987).

42. Teal, T. K., Lies, D. P., Wold, B. J. & Newman, D. K. Spatiometabolic Stratification of Shewanella oneidensis Biofilms. Appl. Environ. Microbiol. 72, 7324–7330 (2006).

43. Murfin, K. E., Chaston, J. & Goodrich-Blair, H. Visualizing Bacteria in Nematodes using Fluorescent Microscopy. J. Vis. Exp. (2012) doi:10.3791/4298.

44. Bao, Y., Lies, D. P., Fu, H. & Roberts, G. P. An improved Tn7-based system for the single-copy insertion of cloned genes into chromosomes of gram-negative bacteria. Gene 109, 167–168 (1991).

45. Trang, K., Bodkhe, R. & Shapira, M. Compost Microcosms as Microbially Diverse, Natural-like Environments for Microbiome Research in Caenorhabditis elegans. J. Vis. Exp. JoVE 10.3791/64393 (2022) doi:10.3791/64393.

46. Shapira, M. & Tan, M.-W. Genetic Analysis of Caenorhabditis elegans Innate Immunity. in Innate Immunity (eds. Ewbank, J. & Vivier, E.) 429–442 (Humana Press, 2008). doi:10.1007/978-1-59745-570-1_25.

47. Bolyen, E. et al. Reproducible, interactive, scalable and extensible microbiome data science using QIIME 2. Nat. Biotechnol. 37, 852–857 (2019).

48. Edgar, R. C. Search and clustering orders of magnitude faster than BLAST. Bioinforma. Oxf. Engl. 26, 2460–1 (2010).

49. Faith, D. P. Genetic diversity and taxonomic priorities for conservation. Biol. Conserv. 68, 69–74 (1994).

50. Shannon, C. E. The mathematical theory of communication. 1963. MD Comput. Comput. Med. Pract. 14, 306–317 (1997).

51. Schindelin, J. et al. Fiji: an open-source platform for biological-image analysis. Nat. Methods 9, 676–682 (2012).

52. Livak, K. J. & Schmittgen, T. D. Analysis of relative gene expression data using real-time quantitative PCR and the 2-ΔΔCT method. Methods (2001) doi:10.1006/meth.2001.1262.

53. Shapira, M. et al. A conserved role for a GATA transcription factor in regulating epithelial innate immune responses. Proc. Natl. Acad. Sci. U. S. A. 103, 14086–14091 (2006).

54. Therneau, T. M., until 2009), T. L. (original S.->R port and R. maintainer, Elizabeth, A. & Cynthia, C. survival: Survival Analysis. (2023).

55. Villanueva, R. A. M. & Chen, Z. J. ggplot2: Elegant Graphics for Data Analysis (2nd ed.). Meas. Interdiscip. Res. Perspect. 17, 160–167 (2019).

